# Behavioral and ERP Markers of Emotion Recognition: Comparing Embodied, Facial, and Emoji Stimuli

**DOI:** 10.64898/2026.07.30.741656

**Authors:** Munna R. Shainy, Arun Sasidharan, Vrinda Marigowda, Prashansa Tripathi, Varsha Vijayan, Advait Basak, Meghna Shekar, Sumit Sharma

**Affiliations:** Axxonet Brain Research Laboratory (ABRL), Axxonet System Technologies Pvt. Ltd., Bengaluru, India; Centre for Consciousness Studies (CCS), Department of Neurophysiology, National Institute of Mental Health and Neurosciences, Bengaluru, India; School of Artificial Intelligence & Data Science (SAIDE), Indian Institute of Technology Jodhpur, Rajasthan, India; Department of Psychology and Counselling, St. Joseph’s University, Bengaluru, India

**Keywords:** emotions, embodied emotions, event-related potential, ERP

## Abstract

While extensive research has been conducted on emotion recognition from facial stimuli, it remains unclear how embodied emotional cues conveyed through body postures, gestures and actions (e.g., stick figures) compare with facial (human faces) and face-like symbolic representations (emoji faces) in shaping behavioral and neural responses. We employed a multimodal approach, combining behavioral measures (accuracy and reaction time) with electroencephalography (EEG) to examine the electrophysiological correlates of emotion recognition. Overall, sixty-six (equal number of males and females; 18–30 year old) participants identified positive, negative, and neutral emotions depicted in the three formats. Behavioral results showed significant effects of both emotion and format on accuracy and reaction time. Emoji faces were recognized with the highest accuracy and fastest reaction times, followed by stick figures and then human faces (p < .001, some comparisons p < .05). Thirty-four participants (17 males and 17 females) underwent EEG, which revealed distinct patterns of event-related potentials (ERPs). The Early Posterior Negativity (EPN) amplitude showed a significant overall effect of format. Post-hoc comparisons indicated that stick figures elicited greater (more negative) EPN amplitudes than human faces during positive emotion recognition (p = .003), and greater amplitudes than both emoji faces (p = .027) and human faces (p = .007) during negative emotion recognition. No significant differences were observed between emoji and human faces. N170 and Late Positive Potential (LPP) amplitudes did not reveal significant differences (all p > .05). Correlation analyses revealed no significant associations between ERP and behavioral measures. Thus, abstract, minimalistic representations like stick figures elicit enhanced early emotion-related processing despite similar early and later processing across formats.

**Highlights:**

- Emotion recognition across realistic (human faces), symbolic (emoji faces) and embodied (stick figures) modalities were compared.
- Emotions were recognized fastest and most accurately in emoji faces format.
- Stick figures format elicited significantly higher EPN compared to other two formats across positive and negative emotions.
- No significant differences were observed in terms of N170 and LPP.
- Embodied, abstract emotional cues modulate automatic emotional appraisal, rather than initial sensory encoding and sustained cognitive evaluation.

## 1 Introduction

According to evolutionary theories, emotions are considered one of the most essential adaptive cues driving our everyday social lives (Hammond, 2006). Emotion is a multifaceted mental state that includes the subjective experience, physiological responses and behavioral or expressive tendencies (Scherer, 2005; Izard, 2010) that are elicited by internal or external stimuli that are relevant to an individual’s goals or wellbeing (Lazarus, 1991; Frijda, 1998) and shaped by both biological predispositions and sociocultural contexts (Mesquita & Walker, 2003; Barrett, 2017). Experimental scientists of diverse fields, including anthropology and sociology, have been studying the intriguing science behind understanding emotions since the 19th century (Thanapattheerakul et al., 2018). Ever since the time of emotion research, the number of emotions expressed, perceived, and identified has increased drastically and so research in this area has exponentially increased, and so has the need to classify them systematically. As early as 1972, Ekman, Friesen, and Ellsworth (1972) identified happiness, excitement, anger, fear, contempt, and sadness as the basic emotions. Later, researchers explored what makes emotions universally distinguishable from each other, especially ones like disgust vs. contempt, where the differences are not that easily apparent (Ekman, 1992). The consensus on classifying emotions into two categories has helped conveniently segregate the vast 66 emotions as basic (anger, anticipation, distrust, fear, happiness, joy, love, sadness, surprise, trust) and secondary emotions (guilt, shame, pride, embarrassment, and jealousy) (Tangney & Dearing, 2002; Tracy & Robins, 2004; Feidakis, Daradoumis & Cabella., 2011). However, researchers consistently consider Russel’s circumplex model of emotions as an efficient standard classification to study emotions, mainly as it further divides emotions based on two simple yet definitive domains: valence (pleasantness/positive or unpleasantness/negative nature of emotions) and arousal (high or low activation caused by the emotions) (Russel, 1980); neither does this system have zero limitations or challenges in assessing emotions (Dzedzickis, Kaklauskas, & Bucinskas., 2020). Notwithstanding the progress in defining and classifying emotions in research, most, if not all, emotions in social contexts depend on having another individual on the opposite side to “recognize” them. Emotion recognition, by definition, is the cognitive ability to perceive and rightly differentiate emotions exhibited by another (Adolphs, 2002). It is to be noted to our awareness, that in scientific literature, emotion “recognition” and emotion “identification” are regarded synonymous, even though when we discuss other concepts in cognitive science, for example, object recognition, we consider recognition as the ability to understand that some-“thing” of a particular class (for example, a fruit) is shown while identification involves differentiation when we narrow down what “exactly” we are shown, i.e., an apple (Beach, 1964; Nosofsky, Clark & Shin, 1989; Thibierge & Morin, 2013). It is empirically confirmed that emotion recognition is a manifestation of the overarching concept of the Theory of Mind (ToM), which refers to the ability to derive inferences about others’ mental states (Frith & Frith, 2005; Aktürk et al., 2020). Emotion recognition and theory of mind are integral to effective social interactions and strengthen close relationships by helping cognize subtle social cues (Philips, Drevets, Rauch & Lane, 2003; Grossmann, 2010; Hoffmann et al., 2010).

### 1.1 Recent Trends in Emotion Recognition Research

In recent decades, researchers have tried to study emotions expressed beyond the traditional form of emotion recognition: facial emotion recognition (FER), which refers to the process of recognizing emotions through facial expressions. However, today, humans have distinct ways of recognizing human emotions from other cues and even inanimate objects and representations (Bowling & Banissy, 2017). Beyond the facial expression, individuals rely on speech features (Dellaert, Polzin, & Waibel, 1996), context (Kang et al., 2021), gestures, and body posture (Stathopoulou & Tsihrintzis, 2011; de Gelder, de Borst & Watson, 2015) to express the emotions and feelings that they experience in social settings (Barbieri et al., 2018; Basharirad & Moradhaseli, 2017; Marrero-Fernández et al., 2014). As technology advances, humans can recognize human-like emotions from written text (Balahur, Hermida, & Montoyo., 2012; Calvo & Mac Kim., 2013; Chatterjee et al., 2019; Sailunaz & Alhajj., 2019), audios (Koolagudi & Rao, 2012), emoticons (Rodrigues et al., 2018), emojis (Duarte, Macedo, & Gonçalo Oliveira, 2019; Völker & Mannheim, 2021; Pfeifer, Armstrong & Lai, 2022), avatars (Sollfrank, et al., 2021), cartoon characters (Schindler et al., 2017) and even in robots (Hsieh, & Cross, 2022). Consequently, significant work has also been done in building software, mobile applications, and neural networking models that can make it easy for one to recognize human emotions in pictures, videos, and texts in multiple languages (Duarte, Macedo, & Oliveira, 2019; Alswaidan & Menai, 2020a) and then convert them into the form of emojis (Duncan, Shine & English, 2016; Alswaidan & Menai, 2020b). Especially in online interactions, emojis like smileys and other symbols are found to aid in a better understanding of emotions from the polysemous text by the sender (Atif, Franzoni, & Milani, 2021). Thus, research on emotion has come a long way, such that several machine learning algorithms, such as deep neural networks, support vector machines, and deep learning, are employed to capture nuanced patterns in EEG signals associated with emotion recognition (Khan & Sharif, 2017; Jafari et al., 2023; Samal & Hashmi, 2024).

A plethora of research has been conducted on recognizing the emotions of humans and animals using a multitude of research methods of varying complexity, from simple survey-based descriptive data to high-end brain imaging techniques (Padhy et al., 2020; Liu et al., 2021; Naga, Marri, & Borreo, 2021). Neuroimaging studies, especially fMRI, established the activation of several neocortical structures, including temporal and medial prefrontal cortices (Jimura, Konishi & Miyashita, 2009) during emotional face recognition. Findings suggest that the fusiform gyrus, amygdala, occipital gyrus, and inferior frontal cortex, during static facial emotion recognition, while the superior temporal sulcus, bilateral inferior occipital cortex, and fusiform gyrus and in general temporal, parietal, and frontal cortices are active during dynamic facial emotion recognition (van de Riet, Grèzes & de Gelder, 2009; Zinchenko, Yaple & Arsalidou, 2018). Studies also have targeted demarcating brain activation during specific emotion recognition; fear signals expressed through the face and body activate the amygdala, superior temporal sulcus, and fusiform gyrus (van de Riet, Grèzes & de Gelder, 2009), happy faces as opposed to sad faces, bilaterally activate anterior cingulate gyrus and amygdala more (Killgore & Yurgelun-Todd, 2004). Electroencephalography (EEG) studies consistently revealed that lateral temporal sites are more active during positive emotion recognition in beta and gamma bands (Zheng, Zhu & Lu, 2017; Brenner et al., 2014). On the other hand, neutral emotions trigger higher alpha responses at parietal and occipital areas, and negative emotions trigger higher delta responses at parietal and occipital areas, and higher gamma responses at prefrontal sites are observed (Zheng, Zhu & Lu, 2017). Further, Gantiva and co-investigators (2020) found that P100 and LPP event-related potentials are more significant during facial emotion recognition, while for emotion recognition in emojis, N170 amplitude was more active. Angry faces elicit more P100 (Smith et al., 2013) and LPP amplitudes than neutral and happy faces (Gantiva et al., 2020).

Most behavioral and psychophysiological studies (both electrophysiological and neuroimaging-based studies) explored facial emotion recognition (FER), and recently, emotions in emojis have been relatively begun to be studied. However, emotions are not limited to human faces or anything symbolic like emoji faces; instead, they could also be expressed via body postures, gestures, and actions, collectively referred to as ‘embodied emotions’ (Prinz, 2008). To our awareness, the difference in behavioural responses and neural markers between emotions expressed via body postures/language compared to emojis and human faces is yet to be studied. Raindel, Liron, and Alon (2018, 2021) have been studying the theory of mind behind body postures in dramatic actions, emotions, and human interactions that they convey, using stick figure configurations depicting a pair of individuals (one neutral stick figure adjacent to another that expresses a dramatic action, for example, compassion, anger, and love). The current study adopted a similar method but used single-stick figure configurations (one stick figure in a picture showing a particular emotion, with no neutral stick figure next to it) to study embodied emotions or emotions expressed via body postures. Thus, the study addresses whether emotion recognition could differ when positive, negative, and neutral emotions are presented in different formats: human faces, emoji faces, and stick figures, behaviorally and neurophysiologically.

### 1.2 Aims of the present study

Thus, the broader objective of this exploratory research study is to examine the differences in recognition of positive, negative, and neutral emotions (specifically happiness, excitement, calmness/serenity, sadness, fear, and anger) when depicted via bodily posture and actions compared to classical human and emoji facial format. First, we examined if differences exist in behavioural responses (accuracy and response time). In a subset of participants, we additionally conducted a simultaneous electroencephalography (EEG) recording to determine the electrophysiological components behind embodied emotion recognition versus during emotion recognition in human and emoji faces formats. Thus, in this study, a multi-pronged approach was adopted to understand the big picture of emotion recognition across the three formats, using both behavioural (reaction time and accuracy) and neural markers (N170, EPN and LPP ERPs) .

## 2 Methods

### 2.1. Participants

In total, 66 (n_EEG recordings_ = 34) young Indian adults (Mean_age_ = 21.97, SD_age_ = 1.78, equal number of males and females) were recruited for this study. Data from two participants were excluded from the analysis due to the poor quality of EEG data.^1^ The participants signed the informed consent form after a proper briefing about the study and the experimental setup. All participants were self-identified right-handed individuals (except one ambidextrous), reported no past or present history of neurological disorder or mental health disorders, and had normal (65.38%) or corrected-to-normal vision. This study was conducted in accordance with the principles embodied in the Declaration of Helsinki and was approved by the internal ethical committee.

### 2.2 Stimuli

The stick figure stimuli set was created for this experiment, inspired by Raindel and co-authors’ (2018, 2021) stick figure configurations. Two images representing three culturally relevant positive (Excitement, Happiness, and Serenity/Calmness), and negative emotions (Afraid/Fear, Anger, and Sadness), and one image representing neutral/no/blank emotions were created. These stimuli were validated and standardized on a separate group of individuals of the same age bracket as that of the study (N=118) (All the images and peer-agreement values can be found here: https://osf.io/xrwb7/?view_only=2a67db8505464398980197e4b15c789a). Similarly, two emoji’s representing the same emotions taken from the open-source Noto Emoji Font set of Google were standardized on another group of individuals (N=112). The percentage of peer agreement was calculated for every corresponding image. Further intercorrelations across all the emoji and stick figure images were analysed to ascertain the possibility of each image depicting more than one emotion. Only images that passed the 80 +/- 2% peer agreement cut-off were finalized (similar to studies involving facial expressions like daSilva, Crager, & Puce 2016; Gantiva, Sotaquirá, Araujo, & Cuervo, 2019). Further, alternative images for certain stick figures that did not meet the cut-off criteria were created, and the same standardization procedure was repeated. The images used in the main experiment all had above 80% peer agreement, and the ones with the least percentage of peer agreement were instead used for practice trials. Similarly, facial stimuli of two models of the Asian-American race (male: 45M and female: 18F) representing each emotion were picked from the NimStim Set of Facial Expressions’ (Tottenham et al., 2009). This was done due to more morphological similarity and familiarity among Indian participants compared to faces from other racial groups, similar to Gantiva, Sotaquirá, Araujo, & Cuervo, 2019’s method. In the EEG experiment, all facial expression images chosen from the NimStim Set of Facial Expressions are converted into monochrome pictures to eliminate the effect of stimulation produced by colour.

### 2.3 Procedure

After signing the consent form, the participants were asked to fill out a form to gather some relevant socio-demographic data. The experiment was hosted via Brain Electro Scan System (B.E.S.S) software. For the experiment, all participants were seated in a sound attenuated room approximately 1.5 meters away in front of the 25.4 x 49.53 cm display monitor. During the experiment, participants were required to fix their focus on the fixation cross (+) placed on the centre of the screen, where all the stimuli were displayed with a white background. The participants were instructed that they would be shown positive, negative, and neutral emotions in three formats: stick figures, emojis, and human faces, and they would have to respond using the three selected keys on the keypad. They responded by pressing the “4” key for negative emotions, “5” key for neutral, and “6” key for positive emotions. Figure 1a depicts typical images used in the study, and the task diagram of the experiment is depicted in Figure 1b. The experiment followed a blocked design, with each block corresponding to a single type of stimulus format (stick figures, emoji faces, or human faces). For the first 20 participants, the blocks were presented in the order of stick figures, emoji faces, and human faces, while the remaining participants completed the blocks in the reverse order in order to counterbalance the stimulus types presentation. There were 3 blocks and each block consisted of 30 trials per emotion category, separated by six 30-second pauses, resulting in a total of 1260 trials. Before the main experiment, all participants were given a 5-trial practice test with seven images (representing each emotion) from each stimulus set, but these images were not used in the main experiment.

**Figure 1a.**
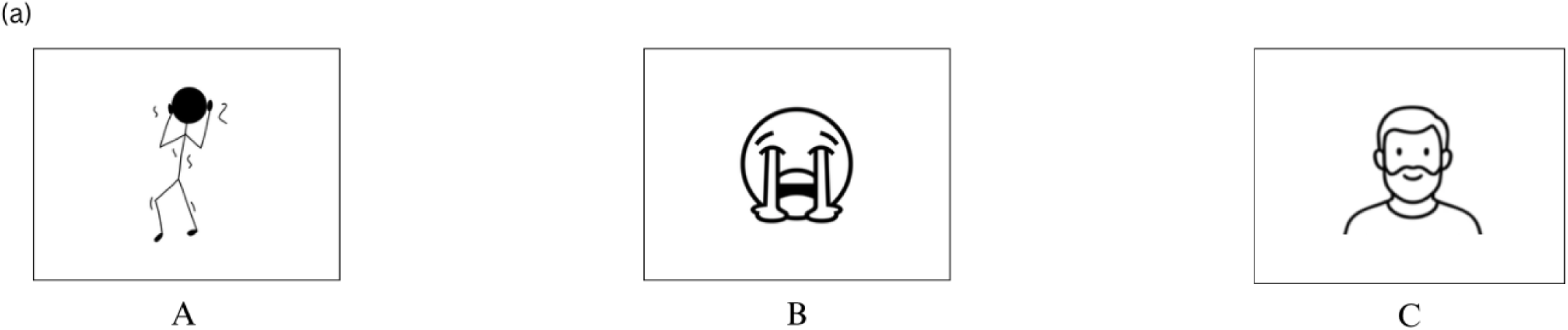
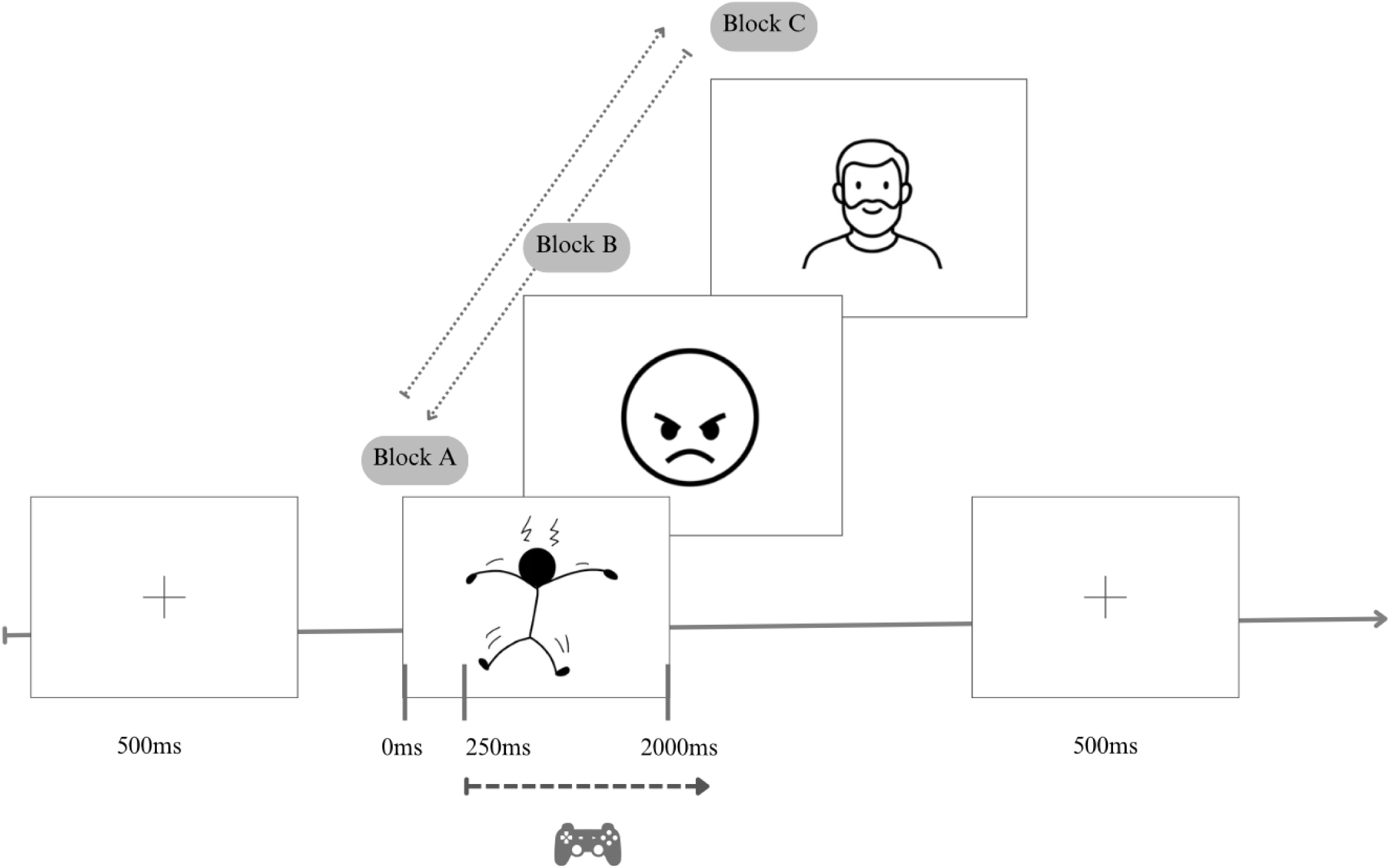
Examples of images depicting (A) stick figures, (B) emoji faces, and (C) human faces used in this study (from Nimstim Set of Facial Expressions). The facial emotion category had one male and female Asian. **Figure 1b.** Trial overview. Images are not to scale

During the EEG-based experiment, a 32-channel EEG cap was attached to the participants after the instructions and a round of practice trial. Half of the participants completed the experimental blocks in the order of stick figures, emoji faces, and human faces, while the other half completed the reverse order; this order was counterbalanced across males and females. In the behavioral experiment, the human faces’ images were in colour and for the participants of the EEG-experiment, the pictures were shown in monochrome as shown below. The room was kept comparatively dimly lit compared to the behavioural experiment. The monitor’s brightness and contrast settings were adjusted to reduce ocular strain during the experimental task.

#### 2.3.1 EEG Recording and Pre-processing

The EEG activity was recorded using a saline-based 32-channel EEG Cap (Axxonet RapidCap). The data was collected from 32 scalp site Ag/AgCl electrodes that followed the standard international 10/20 montage system and the recording was conducted using an EEG-ERP paradigm (BESS FW-32, Axxonet System Technologies Pvt Ltd., Bengaluru) using Brain Electro Scan System (B.E.S.S) software. The EEG signals were digitized with a 1000 Hz sampling rate (0.01-500Hz bandwidth; 24-bits resolution). Along with the EEG data, online keyboard responses and their latencies were recorded simultaneously. For ERP analyses, software bandpass filters of 0.1 Hz to 35 Hz were applied to EEG data, and additionally 50 Hz and 100 Hz notch filters were applied to all EEG recordings as part of preprocessing. The bad channels were semi-automatically detected and verified (n = 5.08 +/- 1.59) in each EEG recording and were interpolated with a spherical spline interpolation technique from all remaining good channel EEG recordings. The data were segmented in epochs from 200 ms before the stimulus onset to 1600 ms after the stimulus onset. Ocular artifact correction was performed based on Principal Component Analysis (PCA) at epoch level. Segments with residual artifacts surpassing voltage steps greater than +/- 100 μV between sample points (0.079 ms to 1300 ms after stimulus onset) were automatically selected and discarded. Further artifacts were then detected and eliminated through visual examination. Subsequently, the same group of emotions within the same format category epochs were clubbed together to form Emotion X Format Categories. (For example, each participant’s respective epochs of images depicting angry, sad, and fearful human faces were averaged together as “Negative X Human Faces”).

### 2.4 Data analysis plan

Statistical analyses of this study were performed mainly in the R programming language via RStudio (R Core Team, 2025). Due to the variability in data distribution across the tested data, tests from robust statistics modality were preferred over traditional inferential statistics for more reliable results.

#### 2.4.1 Behavioural Responses

Accuracy rates and response times of positive, negative, and neutral emotions across three formats are presented respectively in **Figure 2**, created using the *ggplot2* package(Wickham, 2016). Separate Robust ANOVA with trimmed means (ranova, *method = trim*) using *walrus* package (1.0.5 version; 2022) and Yuen’s t test using *DescTools* package (Signorell, 2025), were conducted to evaluate the significance of differences in the time taken for and per cent rate of correct emotion recognition in between formats within each emotion.

**Figure 2.**
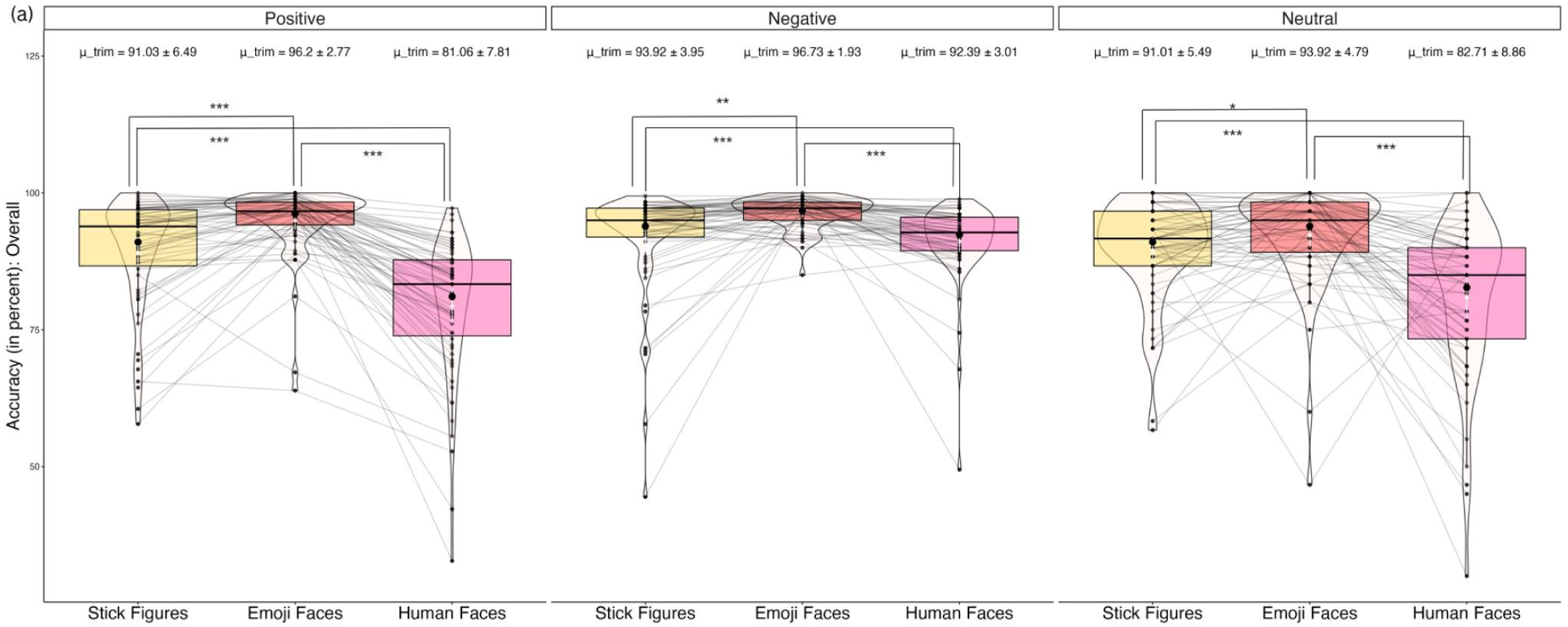
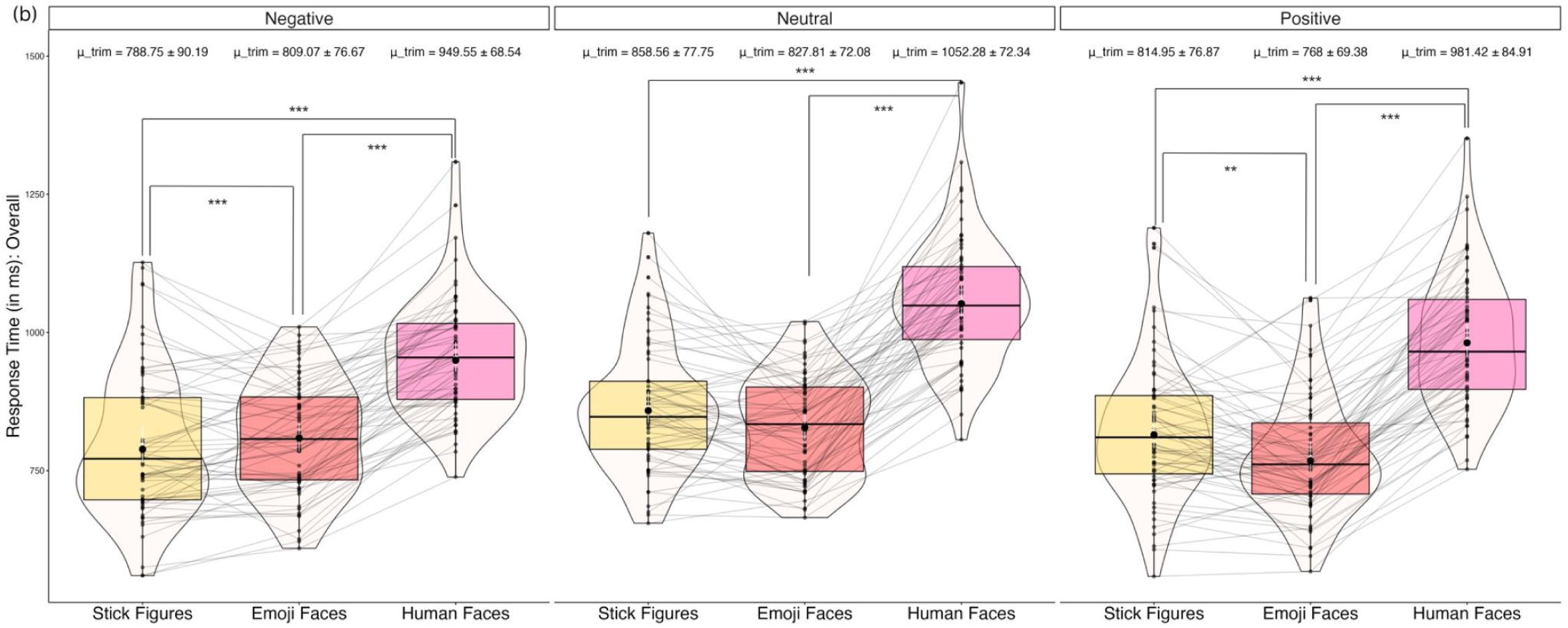
Box plots showing (a) Accuracy Rates and (b) Response Time between Formats (Stick Figures, Emoji Faces and Human Faces) across Emotions (Negative, Neutral and Positive).

#### 2.4.2 Time Domain Analysis: Event-related Potentials (ERPs)

Based on the previous literature, this study considered the mean amplitudes of three ERP components: N170 (face-specific ERP) in lateral occipito-temporal sites, EPN (Early Posterior Negativity) in posterior occipital-parietal sites and LPP (Late Positive Potential) in centro-parietal midline (Bentin et al., 1996; Rossion & Jacques, 2011; Schupp et al., 2003; Hajcak et al., 2010). Additionally, the peak latency of N170 within the time window 80ms to 220ms was also considered (Bentin et al., 1996; Itier & Taylor, 2004). As EPN and LPP are temporally sustained ERP components that last over broader time windows, mean amplitude measures would be more stable and reliable index of neural activity than peak-based measures, which are suited better to temporally discrete components like N170 (Picton et al., 2000; Luck, 2014). The mean amplitude of EPN is considered within the time window of 220 milliseconds (ms) to 500 ms after stimulus onset, and for LPP, the time window of 500 ms to 1200 ms is considered (Schupp et al., 2004b; Schacht & Sommer, 2009; Hajcak et al., 2010; Olofsson et al., 2008). Following manual verification of the grand average (across all formats and emotions) waveform and scalp map distribution, amplitude (N170, EPN, LPP) and latency (N170) data were extracted from only those channels that exhibited these respective ERP components for subsequent statistical analyses as follows (Bentin et al., 1996; Schupp et al., 2003): N170 = P7, P8, TP7, TP8, O1 & O2; EPN = PO3, PO4, O1, O2; LPP = P3, P4, POz, CP3, CP4.

Further, to ensure a fair comparison between negative and positive emotional event-related potentials across the formats, we employed a methodological approach involving the subtraction of amplitudes associated with neutral stimuli from those of negative and positive stimuli across each format. For example,

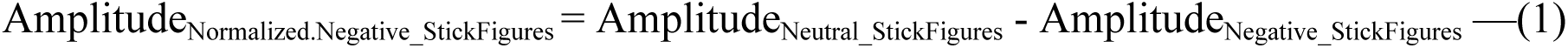

This approach facilitates a nuanced examination of the effects of emotional valence, independent of the image format, by utilizing the respective neutral stimuli as a baseline for comparison. Separate 2 (positive and negative) X 3 (stick figures, emoji faces, human faces) Robust ANOVAs were conducted using *WRS2* package (Mair & Wilcox, 2020), on (1) mean amplitudes (EPN, LPP), (2) peak amplitude of N170, and (3) peak latency period of N170. The Robust ANOVA results, including main effects involving the categories of emotion and format type, and subject as the random factor are reported at a significance level of <0.05. Subsequently, pairwise comparisons across emotion-format comparisons are conducted using separate Yuen’s paired-sample t-tests.

#### 2.4.3 Exploring Relationships between Behavioural and ERP Measures

To answer the question of which neural measure can best explain behavioural responses, accuracy and reaction time, multiple Biweight Midvariance (Rodgers’ robust correlation), correlation tests using *WGCNA* package (Langfelder & Horvath, S, 2008) were carried out with the respective ERP components’ amplitude and latency values.

## 3. Results

### 3.1 Behavioural Responses

Robust Analysis of Variance with 10% trimmed means was conducted to examine the difference in effects of emotion (positive, negative, neutral) and format (stick figures, emoji faces, human faces) on accuracy rate, with the subject as the random factor. The results revealed that emoji faces were recognized with the highest accuracy and fastest reaction times, followed by stick figures and then human faces. Significant main effects of emotion Q(2) = 63.26, p <.001 and format Q(2) = 149.49, p <.001 were found. The interaction effect of emotion and format was also significant, Q(4) = 46.22, p <.001. Three separate Robust one-way repeated measures ANOVA with 10% trimmed means were also conducted to examine the effects of formats on the accuracy rates of each emotion and revealed significant differences in effects across all emotion categories: positive: F(1.58,78.82) = 53.40, p <.001, negative: F(1.53,76.37) = 12.62, p <.001, and neutral: F(1.58,78.82) = 53.40, p<.001.

Similarly, the main effects of emotion Q(2) = 37.13, p <.001 and format Q(2) = 337.78, p <.001 on response time were significant when subjects were considered as a random factor. The interaction effect was also significant Q(4) = 12.69, p=.016. The recognition time was found to differ across each format depicting positive, F(1.81,90.52) = 138.45, p <.001, negative F(1.55,77.57) = 89.20, p<.001, and neutral F(1.84,92.25) = 159.96, p <.001 emotions.

Further, multiple Yuen t-tests were conducted to establish the differences in accuracy rates and reaction time between formats within each emotion category, as presented in Table 1 and Figure 2.

**Table 1.**
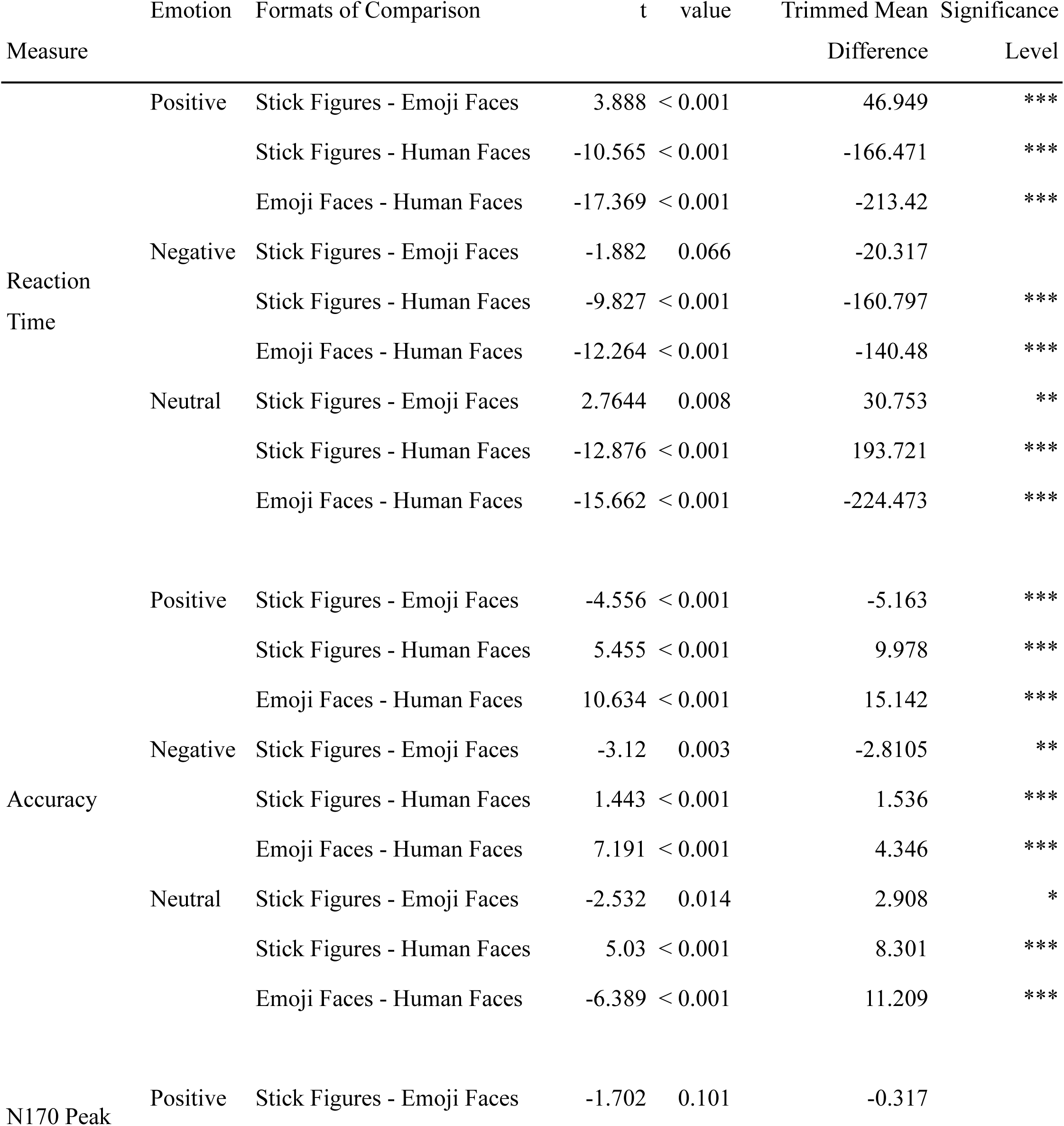
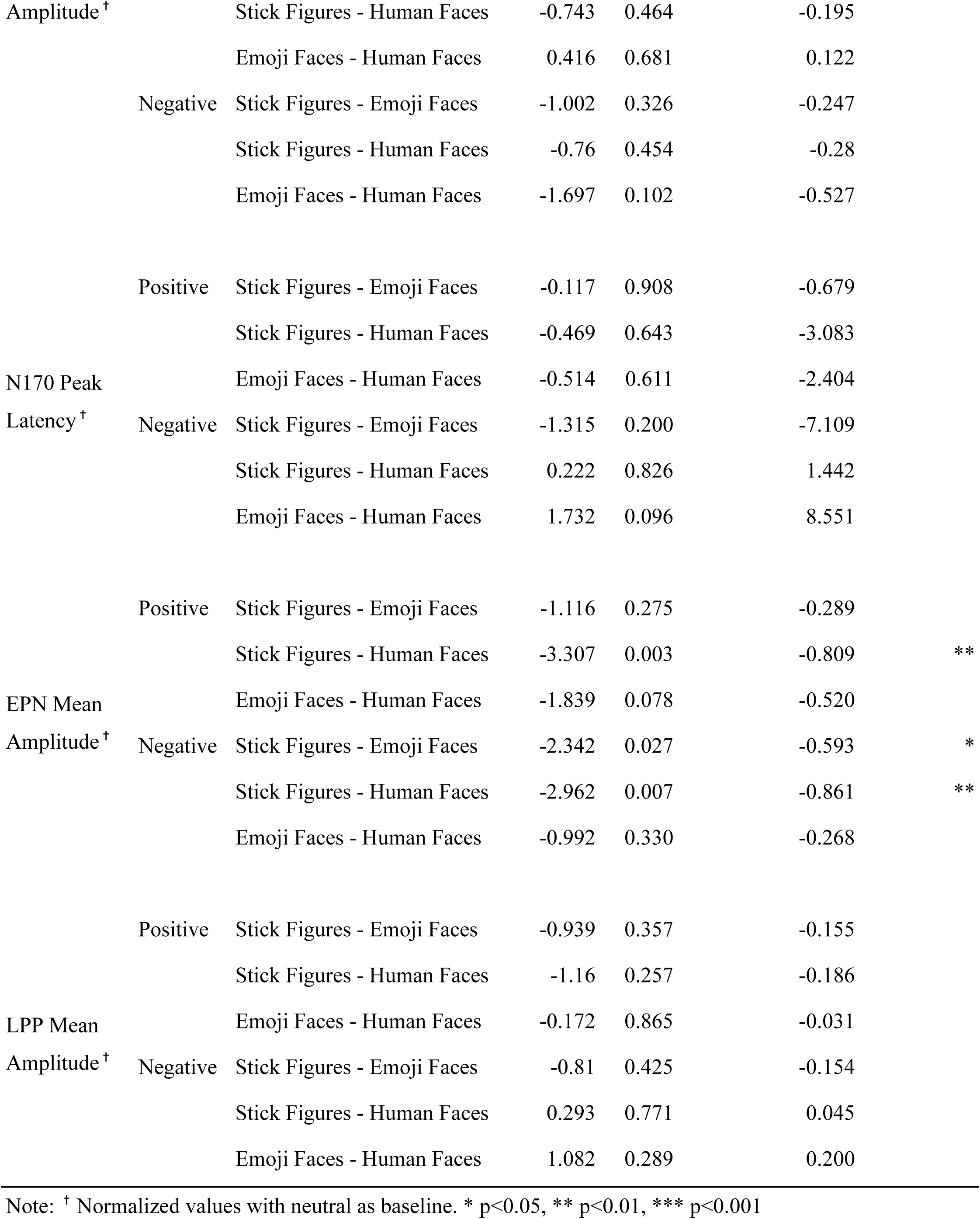
Comparison of Mean Values for Behavioral and Neural Measures Across Different Formats within Each Emotion Category.

### 3.2 Time Domain Analysis: Event-related Potentials

Descriptive statistics for N170, EPN and LPP amplitudes and peak latency of N170 across 2 emotions (positive and negative) and 3 formats (stick figures, emoji faces, human faces) are depicted in **Figure 3** and **Figure 4** depicts the grand average ERP waveforms generated using ERPLAB v12 (Lopez-Calderon, & Luck, 2014) toolbox in MATLAB R2023b and demonstrates the differences in ERP metrics between formats.^2^

**Figure 3.**
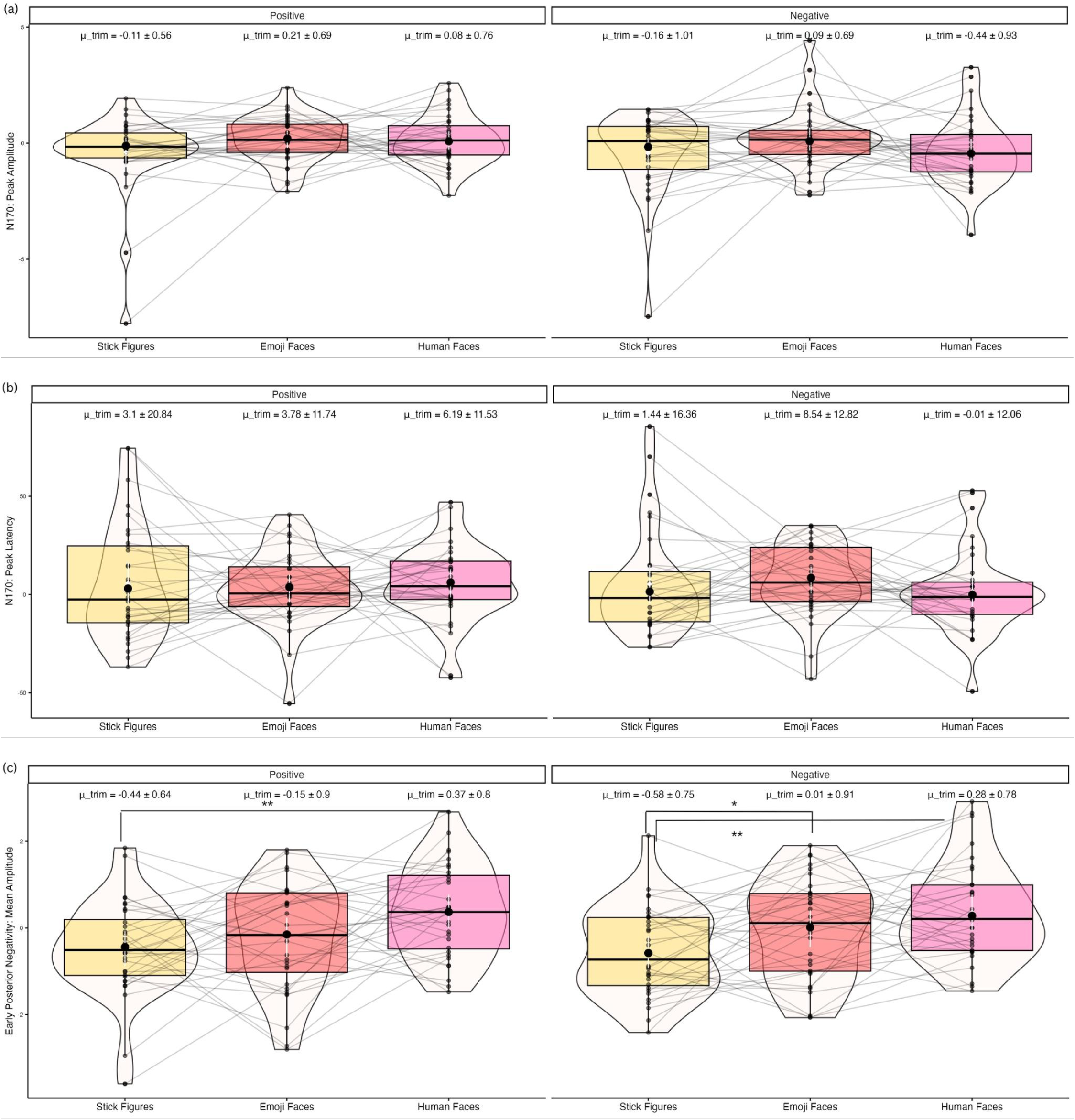
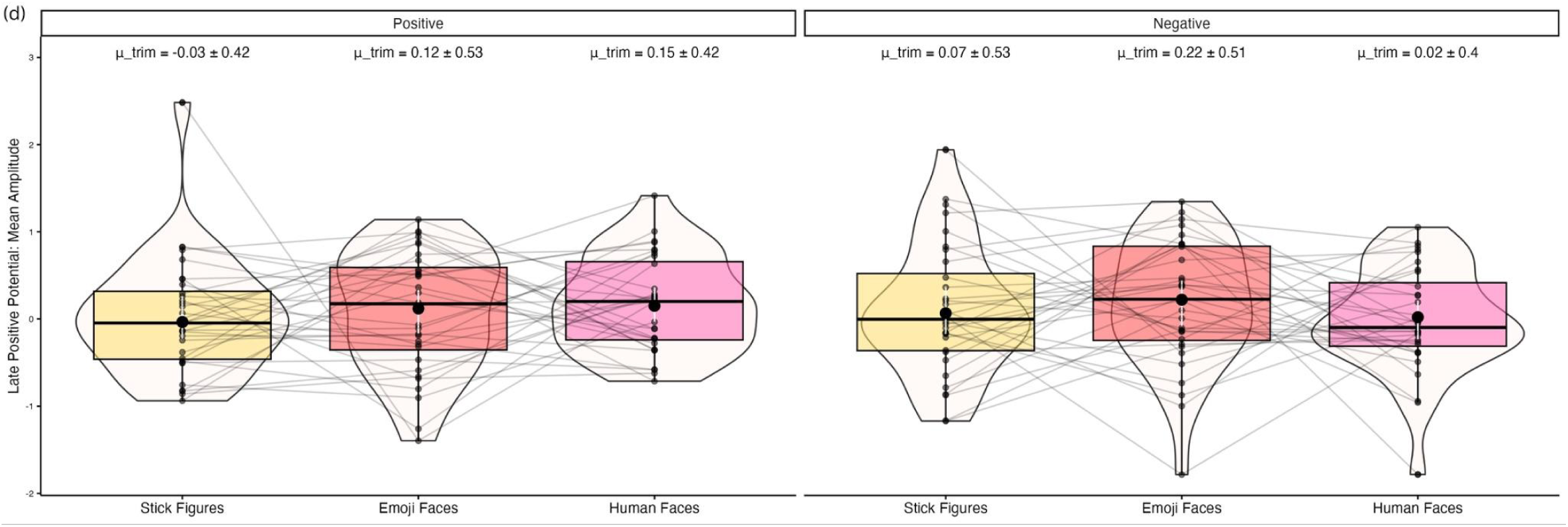
Boxplots Depicting (a) Peak Amplitude and (b) Latency Values of N170 & (c) Mean Amplitude Values of EPN & (d) LPP, relative to that of Neutral images

**Figure 4.**
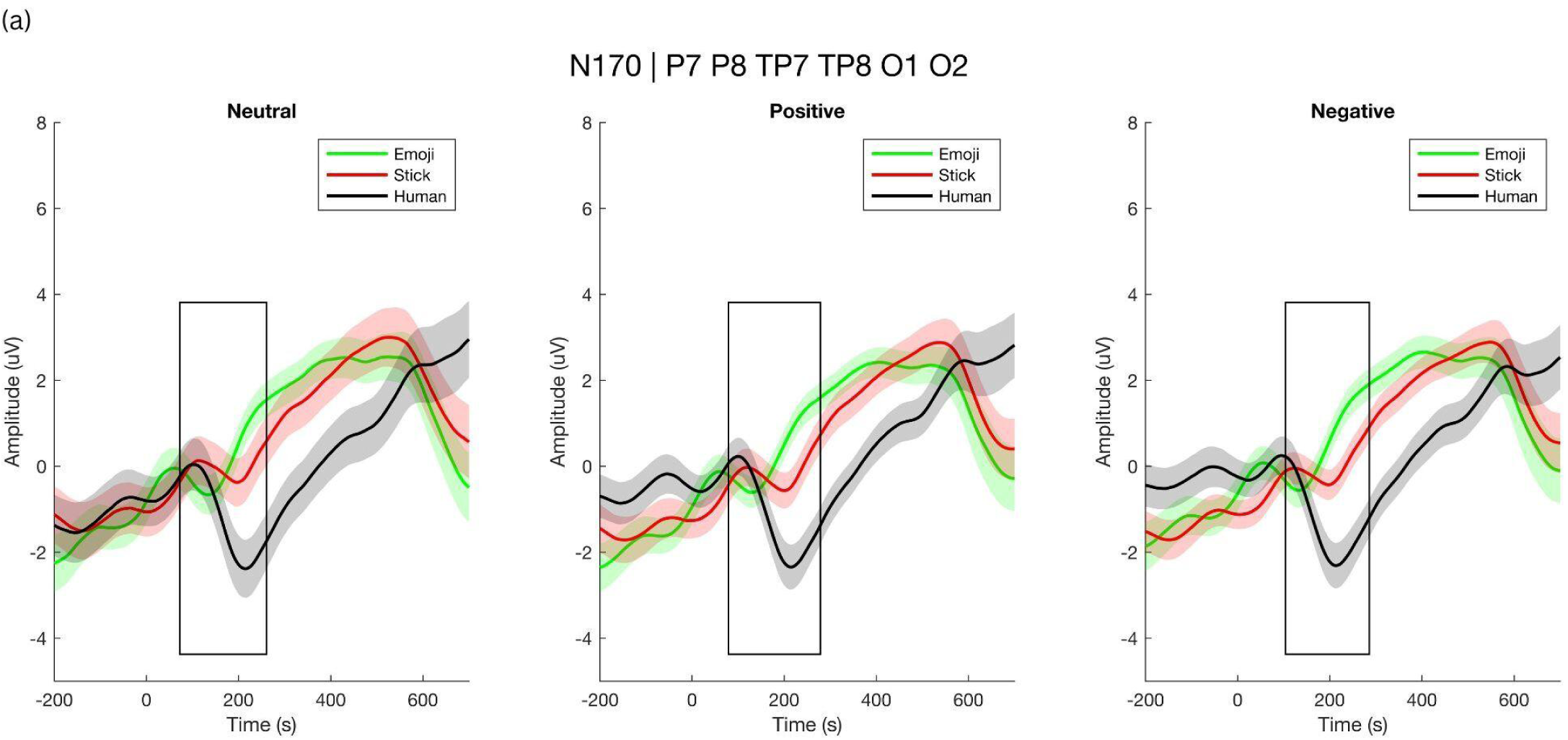
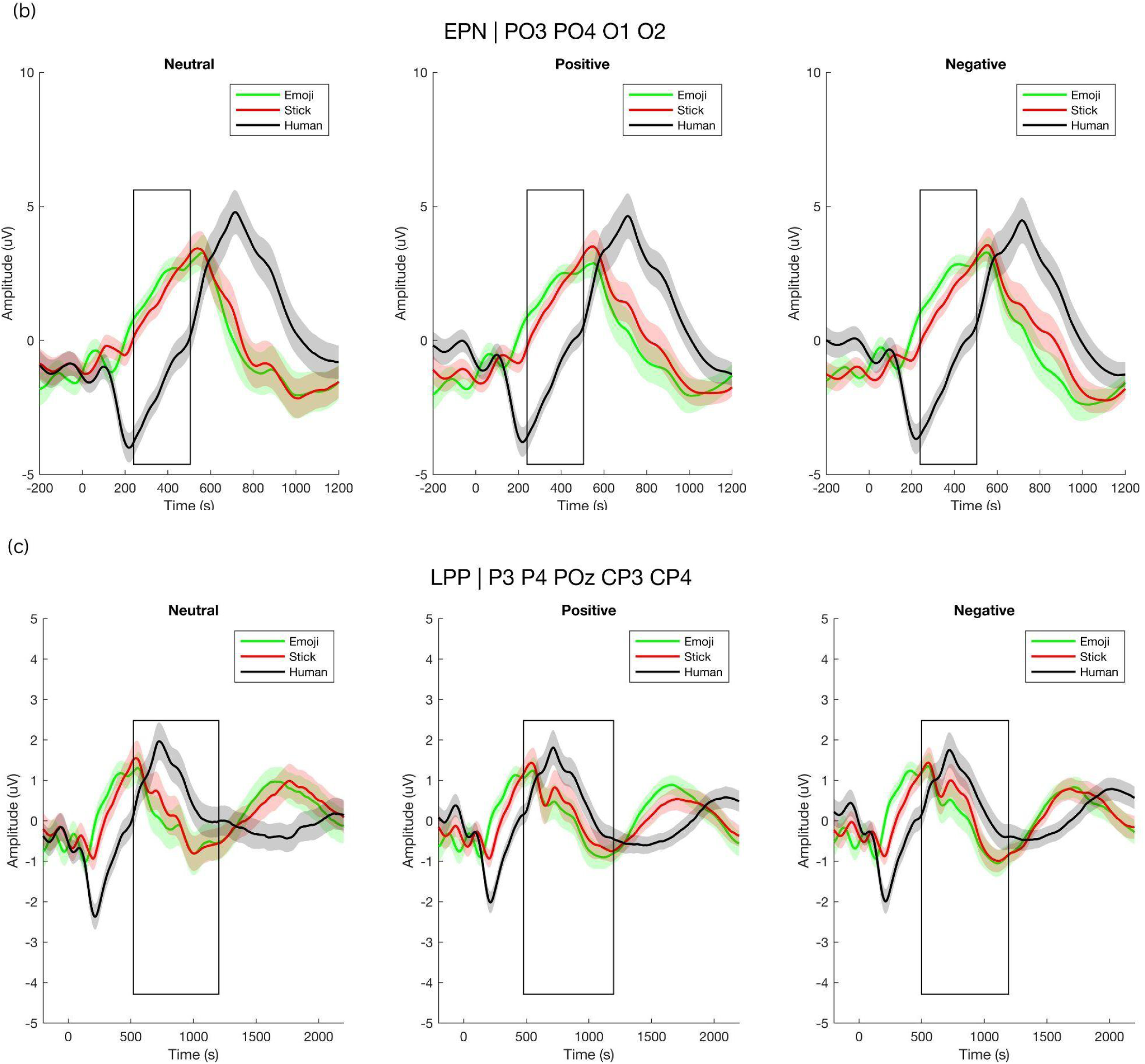
Grand Average Waveforms of (a) N170, (b) EPN and (c) LPP ERP Depicting Differences between Image Formats across Positive, Negative and Neutral Emotions. Shades represent standard error of mean (SEM).

#### 3.2.1 N170

The overall effect of format (Q(2) = 10.67, p = 0.006) was statistically significant, but the overall effect of emotion (Q(2) = 1.91, p = 0.168) and their interaction effect (Q(4) =3.08, p = 0.216) on mean N170 peak amplitudes across P7, P8, TP7, TP8, O1 & O2 channels were not statistically significant upon repeated measures of robust analysis of variance test. Upon testing for format effects within each emotion category, peak N170 amplitudes did not differ across positive (F(50, 2) = 0.019, p = 0.098), but differed across negative emotions (F(45.1, 1.8) = 1.085, p = 0.034). No significant pairwise differences in peak N170 amplitudes were observed during positive emotion recognition between stick figures and emoji faces (t = -1.70, p = 0.101) and stick figures and human faces (t=-0.743, p = 0.464), and emoji faces and human faces (t = 0.416, p = 0.681). Similarly, no significant pairwise differences in peak N170 amplitudes were observed during negative emotion recognition between stick figures and emoji faces (t = -1.00, p = 326), stick figures and human faces (t = 0.760, p = 0.455), and emoji faces and human faces (t =1.70, p = 0.102).

There is no statistically significant effect of format (Q(2) = 0.076, p = 0.963), emotion (Q(2) = 0.024, p = 0.877), or its interaction term (Q(4) = 1.52, p = 0.469), on the N170 peak latency period. Pairwise comparisons also did not result in any significant difference between the effects of formats on N170 peak latency, as presented in Table 1.

#### 3.2.2 Early Posterior Negativity

For EPN mean amplitudes across P7, P8, TP7, TP8, O1 and O2 channels, the main effects of format (Q(2) = 57.71, p = 0.001) but not emotion (Q(2) = 0.0002, p = 0.989) were found significant. The interaction effects were also insignificant (Q(4) = 1.64, p = 0.448). But for format effects within each emotion category, the mean EPN amplitudes varied across positive (F(2,50) = 4.68, p = 0.014), but not negative (F(1.98,49.54) = 2.60, p = 0.085) emotion.

Findings of paired sample Yuen’s t-tests suggested that the amplitude of EPN differed during negative emotion recognition between stick figures and emoji faces (t = -2.34, p = 0.027), stick figures and human faces (t = -2.96, p =0.007), but not for emoji faces and human faces (t = -0.99, p = 0.331). Similarly, EPN during positive emotion recognition in stick figures and human faces (t = -3.31, p = 0.003) were significantly different. However, such a major difference did not exist between emoji faces and human faces (t = -1.84, p = 0.078) and stick figures and emoji faces (t = -1.12, p = 0.275).

#### 3.2.3 Late Positive Potential

No statistically different overall effects were found in terms of the main effects of format (Q(2) = 0.69, p = 0.709), and emotion (Q(2) = 0.000, p = 0.99) upon the mean amplitude of LPP across P3, P4, POz, CP3 and CP4 channels. The interaction effect was also found to be insignificant (Q(4)= 2.07, p = 0.356). Additionally, format effects within the positive (F(2,50) = 1.57, p = 0.217) and negative (F(2,50) = 1.82, p = 0.173) emotion categories were also not significant. Similarly, no pairwise differences in the effects of formats were also found in terms of LPP amplitude during positive or negative emotion recognition, as depicted in Figure 3.

### 3.3 Correlation between Neural Measures and Behavioral Responses

Further, Pearson’s product-moment correlation tests with FDR correction of p-values were employed to identify if there is any relationship between the behavioural responses: accuracy and reaction time and neural measures. The peak amplitude of N170, mean amplitudes of EPN and LPP, and peak latency of N170 ERPs were found to be insignificantly correlated with accuracy and reaction time across all six conditions (Correlation test results of significant ERP and behavioural responses can be found in Supplementary Material 1)

## Discussion

The present multimodal study aimed to identify if embodied emotions (stick figures) are perceived as analogous to facial (human faces) emotions and symbolic (emoji faces) emotions, and which neural marker or markers can best distinguish emotion recognition across these three formats. In terms of accuracy and reaction time, emoji faces were recognized fastest and most accurately, while facial emotions required the most time to be identified and had the least accuracy, suggesting the stick figure format is better than the human faces format but not as best as the emoji format in terms of accuracy and response times. In terms of the neural measures, EPN amplitudes (attentional orientation, emotional arousal, and structural encoding processes) showed differences between the three formats across positive and emotion categories, while no differences in N170 amplitude (lower-order attentional and perceptual processing) and LPP amplitude (higher-order attentional and motivational engagement) were observable.

The behavioural findings indicate a clear effect of stimulus format on emotion recognition, with faster and more accurate recognition in the emoji faces format than stick figures and human faces. This could be explained by the simplified and exaggerated visual features of emojis, which are clear and unambiguous emotional cues leading to rapid perceptual and decisional processes (Churches et al., 2014). However, human faces, irrespective of their ecological validity, have more variability and complexity in emotional expressions, which can, in turn, raise the processing demands and lower accuracy (Calvo & Nummenmaa, 2008). On the other hand, intermediate performance observed in stick figure format can be attributed to their reliance on configural and postural cues rather than fine-grained facial details (de Gelder, 2006; Atkinson et al., 2004), leading to reduced perceptual complexity without fully eliminating ambiguity (Schindler et al., 2017). As a result, they may demand more interpretative processing than emojis, which convey high explicit emotional signals but less than human faces, which involve more subtle and variable features.

The ERP component N170 in the occipito-temporal sites is widely regarded as a face-specific ERP (Krombholz, et al., 2007), (ie; structural encoding of faces) and is also modulated by emotional facial expression (Blau, 2007). A recent study by Gantiva and co-authors (2019) also attributed the increased N170 amplitude to the cortical activation due to structural information (eye and mouth-related features) of emojis compared to that of human faces across both negative and positive emotions. However, in the current study, we observed no such significant difference, despite finding a significant overall effect of format. This suggests that the early structural encoding of formats, indicated by N170 amplitude, is insensitive to variations in formats or emotional valence, and this finding is consistent with prior research (Bentin et al., 1996; Rossion & Jacques, 2011). Stick Figures, due to their abstract simplicity and reliance on minimalistic body-posture cues, require greater visual processing effort and are processed via a ‘different route’ that is not fully reflected in the very early N170 timeframe. This contrasts with human faces (which meet typical face expectations) and emoji faces (which use symbolic/conceptual processing routes). This current finding contradicts Schindler and co-authors’ (2017) finding that N170 is modulated higher for abstract than real faces. In the present study, the ambiguity and deviation of stick figures from typical facial emotion norms may instead impose additional processing demands at later stages of emotion recognition. The stick figure format requires extracting emotional meaning from minimalistic cues, unlike human and emoji faces, where emotions are conveyed through more explicit facial or face-like features. Thus, while human faces may match expectations of a typical face and emoji faces may additionally recruit symbolic or conceptual processing routes, stick figure formats may engage partially distinct interpretive processes beyond the early N170 timeframe.

A pronounced difference in emotional processing is noticeable through the EPN component (a later processing stage around the range of 250-300 ms) (Schupps et al., 2005). The larger EPN amplitude observed in response to both positive and negative stick figures as opposed to the respective emoji and human faces indicates that more allocation of attentional resources and thereby more interaction with the emotional information (Aldunate et al., 2018). This result can be attributed to cortical activity due to the relative novelty of the image and the cognitive load of a more holistic (body posture, angles of body parts, additional cues of space) evaluation of the image to derive the correct emotion depicted. This increased perpetual cognitive load involves inferential reasoning over automatic facial emotion recognition. In this perspective, emoji faces and human faces may trigger more bottom-up attention capture, while stick figures require the integration of contextual and embodied cues to derive emotional meaning, a top-down interpretive mechanism. Thus, an enhanced EPN may be a richer interaction between emotional salience, perpetual ambiguity and structural minimalism.

Although the differences in LPP (later-stage cognitive and motivational processing) did not reach statistical significance, the observed results provide insights into the differences in later-stage evaluation processing of positive and negative emotions across representational formats. The following are just descriptive observations or patterns from our results. The increased LPP amplitudes elicited during positive emotion recognition in human faces, followed by emoji faces and then stick figures, is a pattern that aligns with current literature that indicates the sensitivity of LPP to motivational engagement and the emotional valence of the stimuli (Schupp et al., 2000; Schupp et al., 2004a). In contrast, emoji faces eliciting the highest late positive potential, followed by stick figures and finally human faces, could be related to the affective regulation mechanism (Dennis et al., 2009: Gao et al., 2024) where realistic negative human faces may trigger avoidance responses in later stages of processing while abstract and symbolic emotional formats trigger greater cognitive engagement with comparatively lesser discomfort of raw emotional realism.

## Conclusion

The current study demonstrated both the behavioural and neural markers differences for embodied, facial and symbolic emotion recognition using stick figures, human faces and emoji faces representational formats. It has been shown that the main neural processing differences across emotion recognition across three formats are indicated by the EPN components. Some of the limitations of this study are as follows: (1) no association between individual differences in terms of related traits like emotional reactivity, anxiety, mood experiences and neural activity/behavioral responses were sought (2) valence and arousal ratings of stimulus images were not gathered, (3) results would have been more generalizable if the images used in the study for facial emotions were of Indian faces instead of Asian-American race. Future studies could explore the influence of age, culture and sociodemographics on this emotion recognition task, use high density EEG for increased spatial resolution and robustness of findings and more advanced analyses like intra-brain connectivity, event-related spectral perturbations and localization techniques like sLORETA and beamforming or machine learning algorithms, use wider category of emotions and study how populations with emotion processing deficits (e.g., depression, autism, PTSD) perceive emotions across these formats. However, all the results combined, the study supported the idea that affective processing is modulated by representational image formats.

## Acknowledgments

The authors would like to acknowledge the valuable contributions of Nikita Ghodke, Arjun B. Hargan, and Gauri Mullerpattan to the planning and execution of this project and the staff members of Axxonet System Technologies, Pvt Ltd, for their technical support.

## Funding

This research project was funded by Axxonet System Technologies, Bengaluru,, India

## Availability of data and material

Not available

## Declarations

### Conflict of interest

The authors declare that they have no conflict of interest.

### Consent to participate

All participants provided informed consent prior to participating in the study.

### Consent for publication

All authors consent to publish this manuscript.

### Ethics approval

This study was reviewed and approved by the internal ethical committee at Axxonet Brain Research Laboratory, Axxonet System Technologies. All procedures performed in studies involving human participants were under the ethical standards of the institutional and national research committee and with the 1964 Helsinki declaration and its later amendments or comparable ethical standards. Informed consent was obtained from all individual participants included in the study.

### Declaration of generative AI and AI-assisted technologies in the writing process

During the preparation of this work, the author(s) used ChatGPT to improve the readability of the manuscript in certain complex sentences. After using this tool, the authors reviewed and edited the content as needed and takes full responsibility for the content of the publication.

## Footnotes

1 More information on power analysis estimation can be found in Supplementary Material

2 Figure depicting differences between emotion categories can be found in Supplementary Material

## Notes

### Competing Interest Statement

The authors have declared no competing interest.

