## Supplementary Material for "Behavioral and ERP Markers of Emotion Recognition: Comparing Embodied, Facial, and Emoji Stimuli"

**1.1 Power Analysis**

Power analysis using G*Power 3.1 software (Faul, Erdfelder, Lang, & Buchner, [2007](https://doi.org/10.3758/bf03193146)) was conducted for a priori sample size estimation for analyses related to the primary objectives of the research, that is, behavioral and physiological sub-sections of the study. For the behavioral experiment, the effect size of 0.426, which is almost medium effect as per Cohen’s criteria ([1988](https://doi.org/10.4324/9780203771587)), was set based on the previous literature at the alpha level of 0.05 for repeated measures (3 X 3 groups and measurements), between factors and interactions test. A sample size of 15 was required to fetch a power of 0.80. Similarly, for the neural-markers experiment, the effect size of 0.536 (medium effect according to Cohen’s ([1988](https://doi.org/10.4324/9780203771587))) based on a priori literature at the alpha level of 0.001, requires another n = 21 participants to achieve a power of 0.80.

**1.2.1 Results**

**1.2.1.1 Grand Average Waveforms of N170, EPN and LPP ERP Depicting Differences between Emotion Categories**

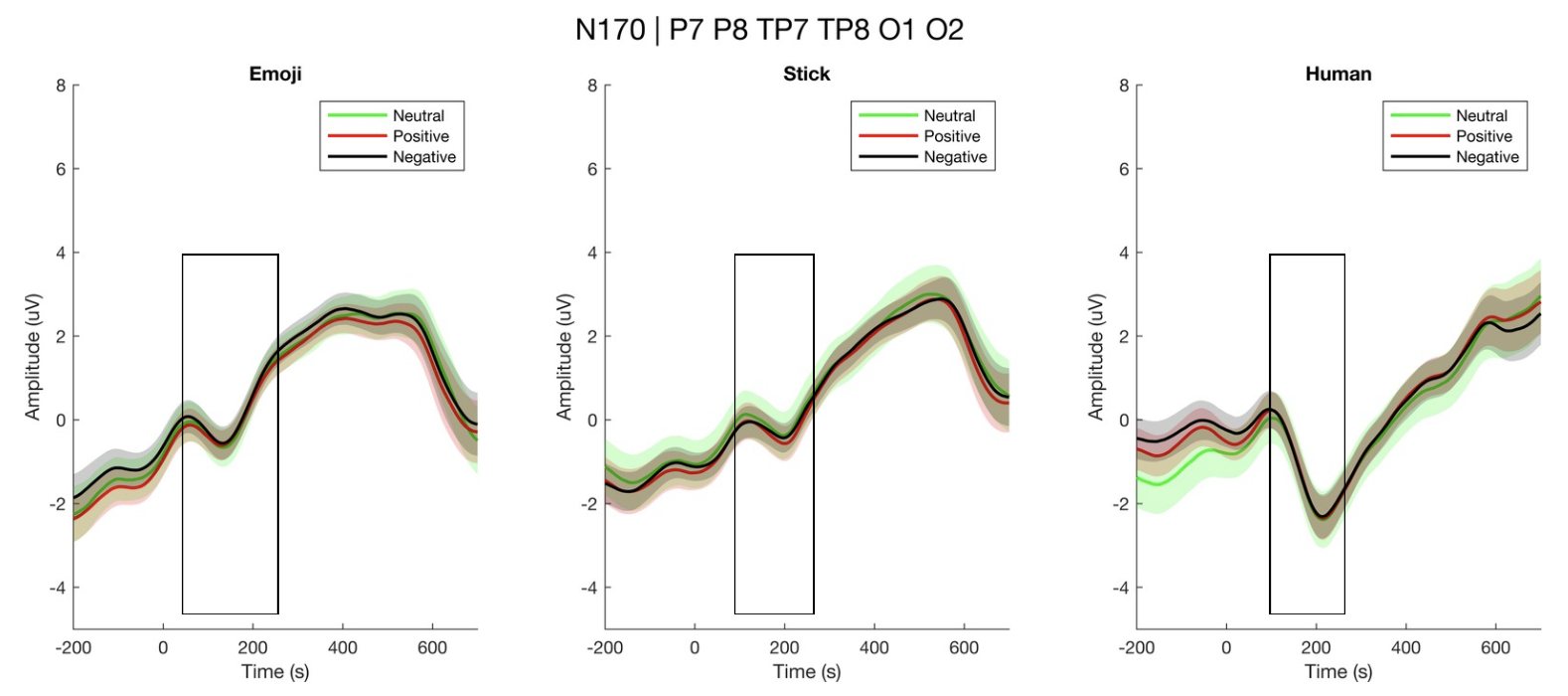

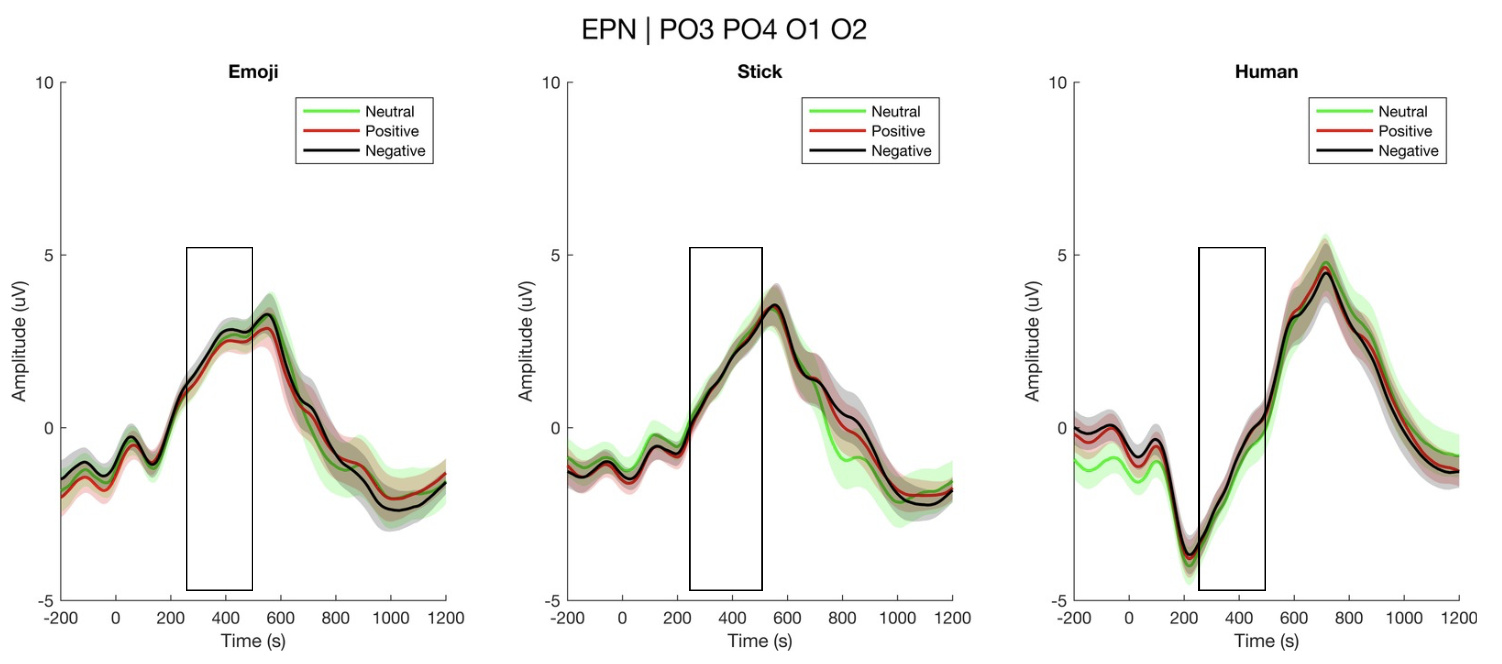

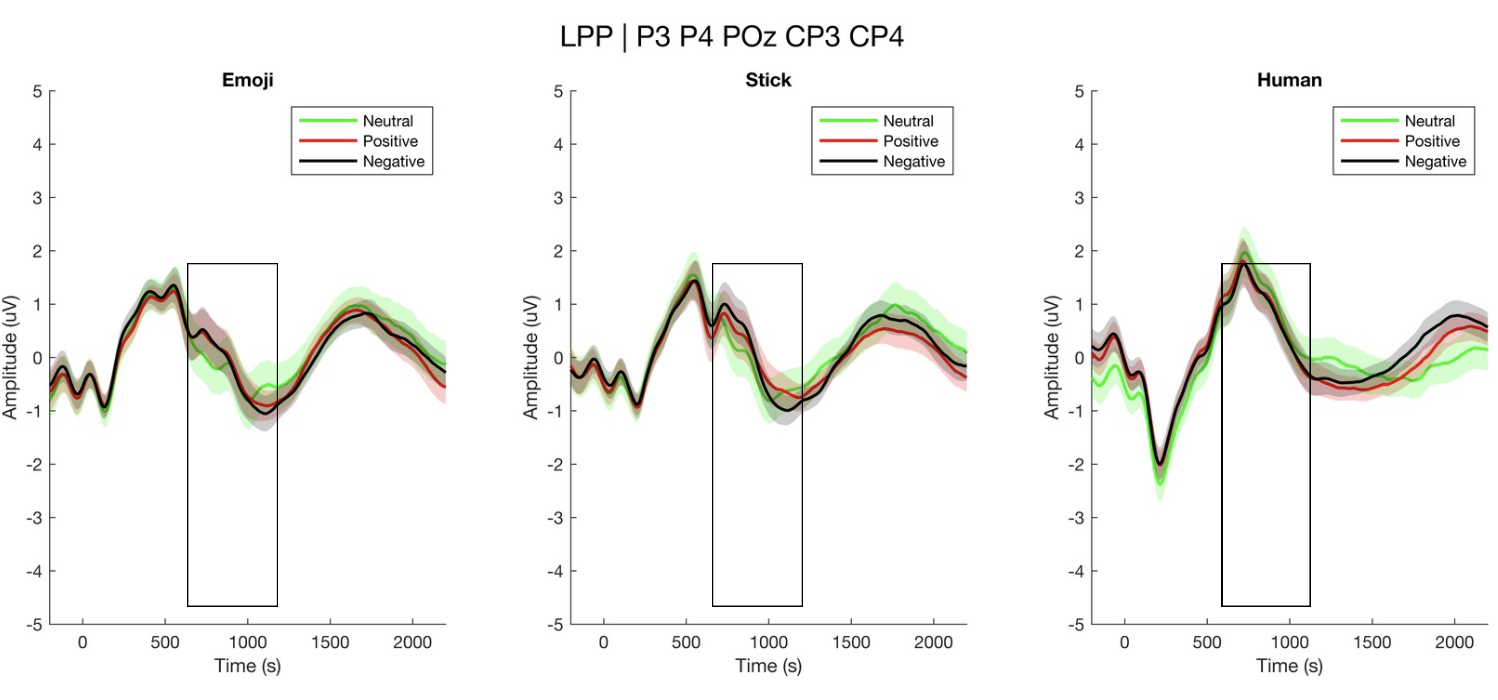

**1.2.1.2 Correlation between ERP and Behavioral Measures**

No significant correlations were found on these tests at any significant levels: 0.05, 0.01 & 0.001.

**Table 1**

*Biweight Midcorrelation (bicor) between ERP Measures and Behavioral Measures across EmotionXFormat*

| Behavioral Measure | ERP Measure | Category | Bicor | *p*-value | FDR-adjusted *p* | Significance |
| --- | --- | --- | --- | --- | --- | --- |
| Reaction Time | EPN | Emoji Negative | 0.1430 | 0.443 | 0.886 |  |
|  |  | Emoji Positive | -0.0187 | 0.920 | 0.970 |  |
|  |  | Human Negative | 0.0069 | 0.970 | 0.970 |  |
|  |  | Human Positive | 0.0160 | 0.932 | 0.970 |  |
|  |  | Stick Negative | 0.3900 | 0.030 | 0.181 |  |
|  |  | Stick Positive | 0.2098 | 0.257 | 0.772 |  |
|  | LPP | Emoji Negative | 0.1199 | 0.521 | 0.965 |  |
|  |  | Emoji Positive | 0.0103 | 0.956 | 0.965 |  |
|  |  | Human Negative | -0.0176 | 0.925 | 0.965 |  |
|  |  | Human Positive | 0.0081 | 0.965 | 0.965 |  |
|  |  | Stick Negative | 0.1534 | 0.410 | 0.965 |  |
|  |  | Stick Positive | 0.1884 | 0.310 | 0.965 |  |
|  | N170 Amplitude | Emoji Negative | 0.1807 | 0.331 | 0.358 |  |
|  |  | Emoji Positive | 0.1710 | 0.358 | 0.358 |  |
|  |  | Human Negative | 0.1835 | 0.323 | 0.358 |  |
|  |  | Human Positive | 0.2091 | 0.259 | 0.358 |  |
|  |  | Stick Negative | 0.3996 | 0.026 | 0.156 |  |
|  |  | Stick Positive | 0.3388 | 0.062 | 0.187 |  |
|  | N170 Latency | Emoji Negative | -0.0802 | 0.668 | 0.796 |  |
|  |  | Emoji Positive | 0.1658 | 0.373 | 0.796 |  |
|  |  | Human Negative | -0.0484 | 0.796 | 0.796 |  |
|  |  | Human Positive | -0.1542 | 0.408 | 0.796 |  |
|  |  | Stick Negative | 0.1003 | 0.591 | 0.796 |  |
|  |  | Stick Positive | -0.2271 | 0.219 | 0.796 |  |
| Accuracy | EPN | Emoji Negative | -0.0639 | 0.733 | 0.923 |  |
|  |  | Emoji Positive | 0.2398 | 0.194 | 0.630 |  |
|  |  | Human Negative | -0.0318 | 0.865 | 0.923 |  |
|  |  | Human Positive | 0.0831 | 0.657 | 0.923 |  |
|  |  | Stick Negative | 0.0182 | 0.923 | 0.923 |  |
|  |  | Stick Positive | -0.2315 | 0.210 | 0.630 |  |
|  | LPP | Emoji Negative | 0.0496 | 0.791 | 0.796 |  |
|  |  | Emoji Positive | 0.3167 | 0.083 | 0.496 |  |
|  |  | Human Negative | 0.0485 | 0.796 | 0.796 |  |
|  |  | Human Positive | -0.1545 | 0.407 | 0.646 |  |
|  |  | Stick Negative | 0.2110 | 0.255 | 0.646 |  |
|  |  | Stick Positive | -0.1469 | 0.430 | 0.646 |  |
|  | N170 Amplitude | Emoji Negative | -0.2479 | 0.179 | 0.536 |  |
|  |  | Emoji Positive | 0.0989 | 0.596 | 0.690 |  |
|  |  | Human Negative | -0.1524 | 0.413 | 0.690 |  |
|  |  | Human Positive | -0.1059 | 0.571 | 0.690 |  |
|  |  | Stick Negative | -0.0746 | 0.690 | 0.690 |  |
|  |  | Stick Positive | -0.3965 | 0.027 | 0.163 |  |
|  | N170 Latency | Emoji Negative | -0.2531 | 0.170 | 0.509 |  |
|  |  | Emoji Positive | -0.3157 | 0.084 | 0.502 |  |
|  |  | Human Negative | 0.0693 | 0.711 | 0.711 |  |
|  |  | Human Positive | -0.1216 | 0.515 | 0.711 |  |
|  |  | Stick Negative | -0.0943 | 0.614 | 0.711 |  |
|  |  | Stick Positive | 0.1330 | 0.476 | 0.711 |  |
